# cuBayes: GPU accelerated FreeBayes that achieves 1-minute whole-genome SNV calling while maintaining algorithmic semantics

**DOI:** 10.64898/2026.06.12.731910

**Authors:** Anders Pitman, Cathy Yang, Yi Qiao

**Affiliations:** Department of Human Genetics, University of Utah, Salt Lake City, UT, USA; Department of Biomedical Informatics, University of Utah, Salt Lake City, UT, USA

## Abstract

Next-generation sequencing now produces whole-genome data in hours, but downstream variant calling remains a multi-hour to multi-day bottleneck that excludes genomic analysis from time-critical clinical settings. GPU acceleration offers a natural path forward — variant calling is inherently parallelizable across genomic positions — yet open-source infrastructure for porting existing algorithms to GPU hardware remains limited, leaving many widely-used tools without accelerated implementations. FreeBayes, a haplotype-based variant caller central to the 1000 Genomes Project and to multi-sample tumor evolution analyses, exemplifies this gap: it is natively single-threaded despite its algorithmic suitability for parallelization. We present cuBayes, a CUDA implementation of FreeBayes germline SNV calling that completes HG002 and HG004 2×250bp Illumina 60× whole-genome analysis in one minute (as opposed to hours if not days with manual region-based CPU parallelization) on a single NVIDIA RTX 6000 Ada GPU, while producing variant calls with 99.97% concordance to the CPU reference. cuBayes is structured around an atom/molecule architecture in which reusable functional units (BAM decompression, position-wise pileup, batch coordination) are cleanly separated from algorithm-specific logic, providing a foundation intended to support acceleration of additional sequence analysis algorithms without redundant low-level engineering.

## INTRODUCTION

High-throughput DNA and RNA sequencing have become foundational tools across biomedical research and clinical care, with applications spanning nearly every research sub-field and clinical conditions. A particular category of conditions, however, stands to benefit most from sequencing-based approaches and yet remains underserved: those for which clinical decisions cannot wait. Critically ill neonates, acute presentations of rare diseases, and aggressive cancers including pediatric brain tumors and adult glioblastoma share a common feature — genomic insights are most actionable when delivered within hours to days of sample acquisition. The clinical value of rapid turnaround has been repeatedly demonstrated: rapid whole-genome sequencing for critically ill neonates was first reported at ∼50 hours in 2012^1^, reduced to under 20 hours by 2019^2^, achieved in 7 hours 18 minutes via nanopore sequencing in 2022^3^, and most recently completed in under 4 hours using prototype sequencing-by-expansion technology^4^. These efforts validate the clinical impact of rapid genomics but also expose its central limitation: each required prototype hardware, hand-optimized workflows, and dedicated multidisciplinary teams — a level of resource intensity unavailable in typical clinical or research settings.

Sequencing itself is no longer the rate-limiting step in most modern workflows; downstream computational analysis is^5^. The most widely used sequence aligners (e.g. BWA MEM^6^) and variant callers (e.g. the GATK best-practices pipeline^7^ and FreeBayes^8^) were developed during the 1000 Genomes Project era^9^ for compute environments characterized by slow storage, modest core counts, and the absence of hardware accelerators. As a result, even on contemporary hardware, the GATK best-practices pipeline routinely requires tens of hours for a single 30× whole genome and approaches 24 hours at 50× coverage on a dedicated HPC node^10^, while FreeBayes is natively single-threaded, and even with manual, region-based parallelization, still requires hours for a 60× whole genome despite its algorithmic suitability for parallelization across genomic positions. Hardware acceleration has emerged as a natural response. Commercial platforms (NVIDIA Parabricks, Illumina DRAGEN) have demonstrated order-of-magnitude speedups for GATK-style germline workflows^11^, and academic work has accelerated variant calling components (e.g. the Pair-HMM kernel central to GATK HaplotypeCaller^12^) and alternative approaches such as the deep-learning-based DeepVariant^13^. Together, these efforts establish that hardware (or specifically, Graphics Processing Unit, GPU) acceleration of variant calling is technically feasible and that meaningful performance gains are achievable.

What remains absent is the open infrastructure that would translate this feasibility into broad adoption. Two barriers persist. First, GPU programming requires expertise distinct from algorithm design — host–device coordination, memory hierarchies, kernel orchestration — that most biomedical method developers do not have. Second, the existing acceleration landscape offers no foundation to build on: commercial platforms are closed-source and vendor-locked, while academic efforts are typically tool-specific, with bespoke architectures designed in isolation that limit reuse. A method developer wishing to accelerate a new algorithm must currently re-implement low-level I/O, decompression, memory management, and parallel coordination from scratch. In our prior work on quickBAM^14^, we demonstrated that common bioinformatics operations such as BAM file decompression and pileup computation can be effectively parallelized, and we packaged the underlying primitives as a reusable software development kit targeting multi-core CPU architectures. Reimplementing existing tools on top of quickBAM, including samtools flagstats and the snp-pileup utility of FACETS, yielded order-of-magnitude speedups without tool-specific re-engineering of low-level data access. Extending this approach to massively parallel GPU architectures — where the throughput potential is substantially greater but the engineering complexity is correspondingly higher — is the natural next step toward making genomic analysis even more accelerated while maintaining broad developer accessibility.

Here we present cuBayes, a CUDA-based, GPU-accelerated implementation of FreeBayes germline SNV calling that completes a 60× whole-genome analysis in one minute on a single NVIDIA RTX 6000 Ada GPU while preserving algorithmic equivalence with the reference implementation. FreeBayes is a haplotype-based variant caller that contributed to the 1000 Genomes Project and natively supports joint calling across samples — capabilities heavily used not only in population genetics but also in multi-sample tumor evolution analyses^15^ — yet has no existing GPU implementation. Beyond the cuBayes implementation itself, we organize its design around a two-level abstraction intended to generalize to other algorithms. At the **atom** level, we implement indivisible functional units corresponding to basic GPU-programming and bioinformatics primitives — host-to-device buffer transfers, BGZF block decompression, position-wise pileup kernels, genotype evaluation kernels — each with well-defined interfaces. At the **molecule** level, we compose atoms into self-contained functional units that handle complete subtasks within a sequence analysis workflow — BAM decompression with partitioning, pileup with cross-batch handling, genotype calling with VCF emission — exposing simple semantic interfaces to higher-level workflow code. The result is an implementation that achieves sub-minute whole-genome SNV calling on commodity GPU hardware while providing a concrete, reusable pattern for accelerating additional algorithms. We provide cuBayes as a containerized release for immediate testing. Source code will be released under a permissive open-source license alongside journal publication.

## Results

We implemented cuBayes, a CUDA-based GPU-accelerated implementation of germline SNV calling that is mathematically equivalent to FreeBayes v1.2.0 SNV-only semantics. We first benchmarked its performance and assessed call concordance against the reference FreeBayes implementation using two GIAB benchmark samples, HG002 (NA24385) and HG004 (NA24143), sequenced as 2×250 bp Illumina at 60× coverage with novoalign-aligned BAM files obtained from the GIAB data release at GRCh38, then extended this comparison to other GIAB samples (HG001-HG007). Beyond the cuBayes implementation itself, we designed a two-level abstraction architecture that encapsulates reusable software components and common GPU acceleration patterns, intended to support the acceleration of additional variant-calling algorithms with reduced engineering effort.

### Sub-minute whole-genome SNV calling on commodity GPU hardware

We benchmarked cuBayes against the reference FreeBayes implementation, run with manual region-based parallelization across non-overlapping genomic regions on multi-core CPU. All runs used GIAB benchmark samples HG002 and HG004 as Illumina 2×250 bp WGS data at 60× coverage, with novoalign-aligned BAM files retrieved from the GIAB data release at GRCh38^16^. All measurements were performed on a workstation equipped with an AMD Ryzen Threadripper 7975WX (32 cores /64 threads) and a single NVIDIA RTX 6000 Ada GPU (48 GB). Detailed hardware and software configurations are provided in **Methods**.

cuBayes completed end-to-end germline SNV calling on HG002 in **68 seconds** and HG004 in **74 seconds** (**Figure 1A**). The reference FreeBayes implementation, run with manual region-based parallelization across non-overlapping genomic windows on the same workstation, required **5 hours 45 minutes at 12 threads** and **2 hours 22 minutes at 32 threads** for HG002**; and 5 hours 32 minutes at 12 threads** and **2 hours 18 minutes at 32 threads** for HG004. Increasing thread count from 12 to 32 yielded a 2.3-fold reduction in runtime, and further CPU-scaling likely would require multi-node, distributed computing with additional overhead in realistic settings. cuBayes on a single GPU therefore outperforms manually parallelized CPU FreeBayes by more than 125-fold on hardware available within a single commodity workstation.

**Figure 1.**
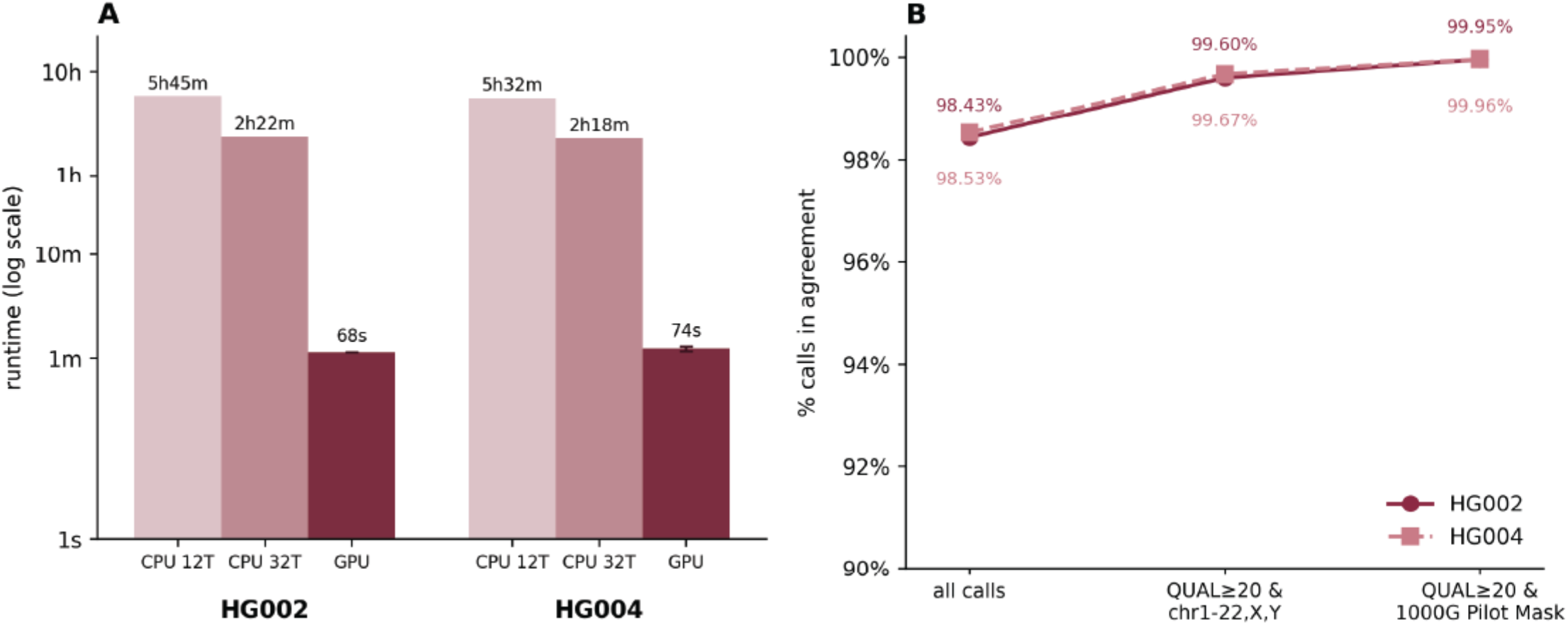
cuBayes runtime and concordance performance using HG002 and HG004. **A)** cuBayes runtime performance compared to the stock version of FreeBayes at 12 threads and 32 threads. The Y axis is in log scale. **B)** cuBayes variant calling concordance compared with the stock FreeBayes at three comparison levels – all calls, high quality variants, and high quality variants within 1000 Genome published accessible regions.

We assessed call equivalence by comparing cuBayes output VCFs against the reference FreeBayes implementation on the same input, using hap.py^17^ for concordance metrics. Concordance is measured at three levels: level 1 is comparing all calls straight from the two callers; level 2 is restricting the comparison to main chromosomes (chr1-22,X,Y) and filter for quality score >= 20 variants; and level 3 is further restricting level 2 variants to high confidence genomic regions as published by the 1000 Genome accessibility map^9^. On HG002, cuBayes and stock FreeBayes results achieved 98.43%, 99.6%, and 99.95% concordance at the three levels respectively; and on HG004, 98.53%, 99.67%, and 99.96% respectively (**Figure 1B**). We further extended this analysis to other Genome In A Bottle samples that represent different sequencing configurations and depths (**Figure 2**).

**Figure 2.**
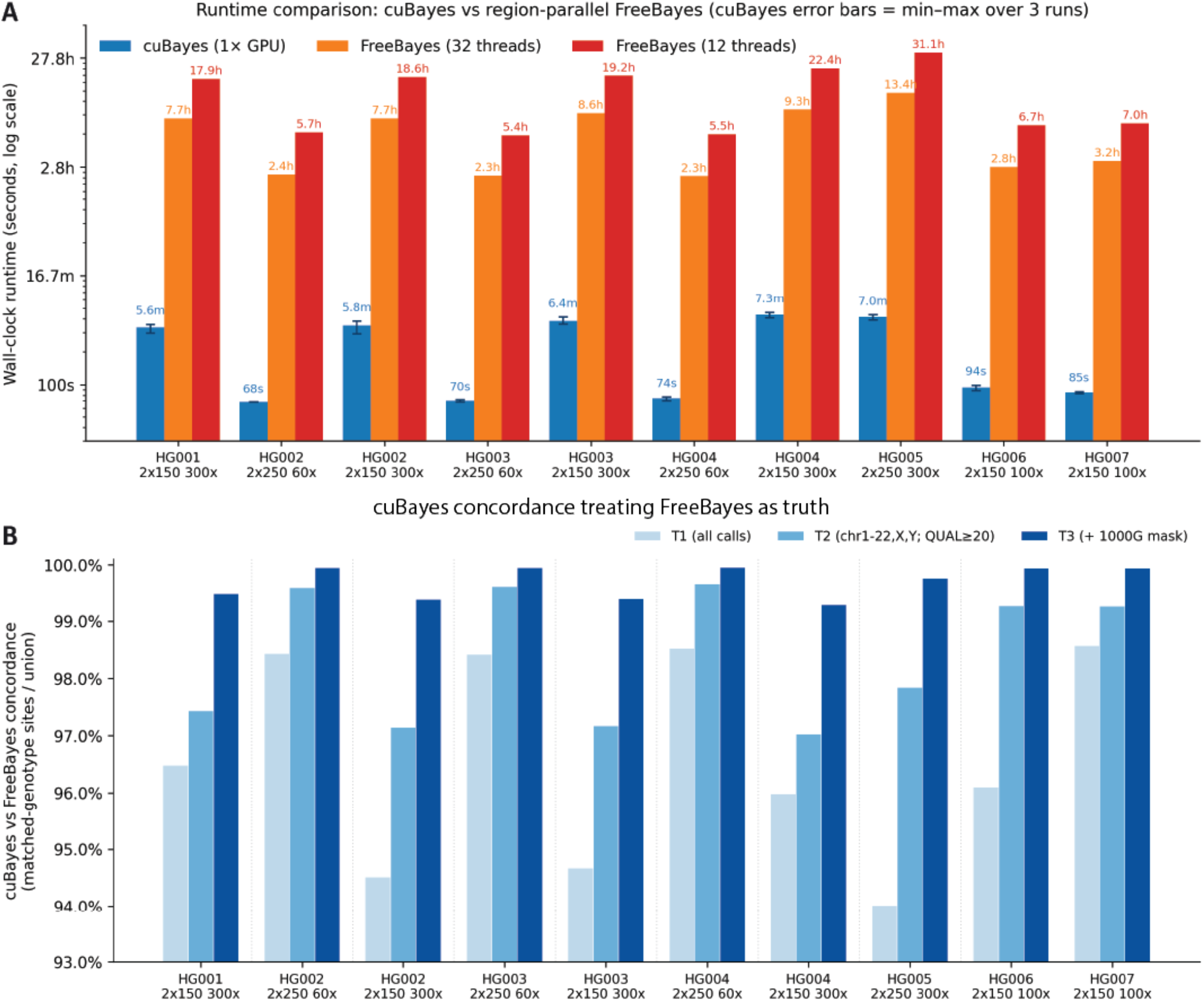
cuBayes vs FreeBayes comparison across HG001-HG007. The sequencing specifications are labeled under the sample names. **A)** runtime comparison. cuBayes were run 3 times per analysis to evaluate variance. cuBayes runtime generally correlates with sequencing depth. **B)** concordance analysis. Tier 1 (T1) compares all calls as produced without any filtering; Tier 2 filters on variant quality and restrict to main chromosomes; and Tier 3 restricts the analysis further to the published high quality genome regions (1000 genome pilot-style mask)

### An atom/molecule architecture provides reusable infrastructure for accelerated variant calling

We designed cuBayes from the ground up with code reusability in mind. To enable the acceleration of not only FreeBayes, but other sequence analysis algorithms, we developed a set of supporting infrastructure that handles BAM access, decompression, partitioning, and pileup construction. We deliberately separated these concerns into a two-level architecture intended to make the supporting infrastructure reusable easily (**Figure 3**).

**Figure 3.**
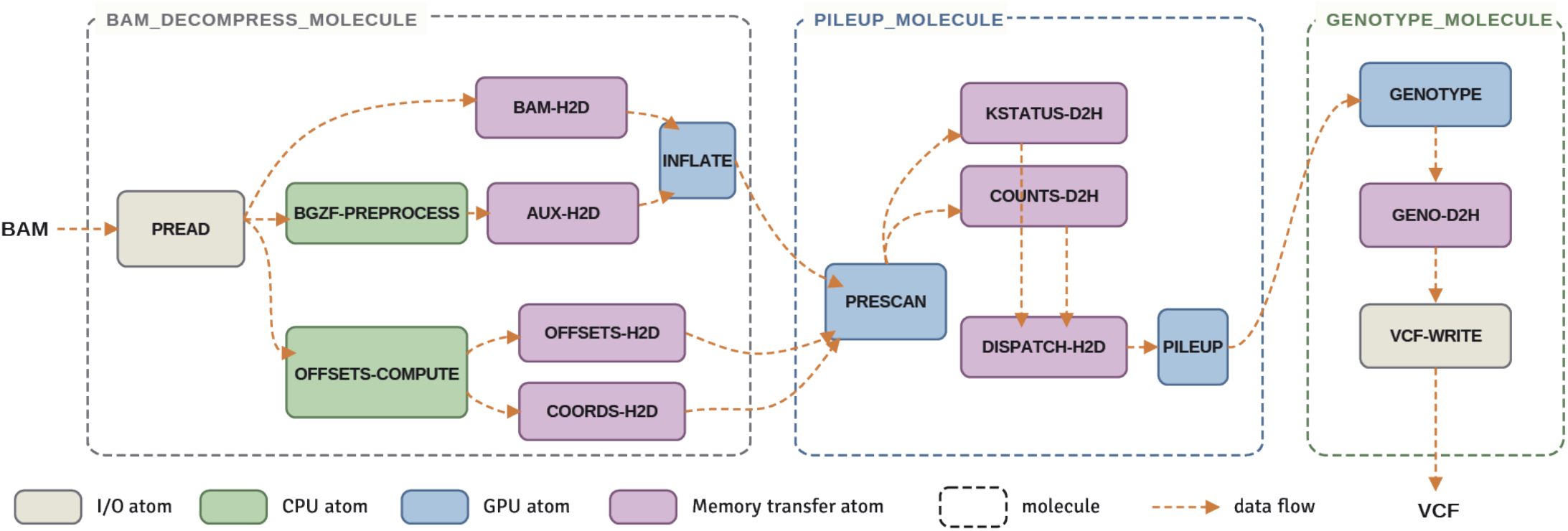
Atom and Molecule architecture implemented in cuBayes. Each atom is a self-contained, indivisible unit of function that can execute independently from other atoms. When they are chained together, they form a data path which defines data dependency and excerpts back pressure to allow proper synchronization while permitting independent atoms to execute in parallel. These atoms and molecules can be reused to compose a different algorithm other than FreeBayes, with much of the code unchanged. For example, all the *-D2H atoms are instantiations of the same D2H (device to host memory transfer) atoms with different source and destination buffer pointers.

**Atoms** are indivisible functional units that encapsulate a single GPU-programming or bioinformatics primitive. cuBayes implements a set of foundational atoms including paged file reads (PREAD), host-to-device and device-to-host memory transfers (H2D, D2H), BGZF block decompression (INFLATE), block-boundary scanning to identify alignment record offsets (PRESCAN), position-wise pileup construction (PILEUP), and Bayesian genotype evaluation (GENOTYPE). Each atom exposes a narrow interface defined entirely by its input and output buffers, and can be invoked independently or composed into larger units.

**Molecules** are self-contained functional units assembled from atoms that handle complete subtasks within a sequence-analysis workflow. cuBayes implements three top-level molecules: a BAM decompression molecule that streams compressed alignment data from storage, partitions it across 16 kb genomic windows, and produces raw BAM records; a pileup molecule that aggregates aligned read evidence position-wise across a batch of partitioned regions; and a genotype molecule that calculate posterior probabilities for candidate variants under FreeBayes’ haplotype-aware model and emits VCF records. Molecules expose semantic interfaces — for example, “decompress and partition this BAM region” — that hide host-device coordination, batch sizing, and memory management from calling code.

## DISCUSSION

### Comprehensive whole-genome variant calling can be substantially accelerated using existing commodity hardware

cuBayes demonstrates that a foundational genomic analysis — whole genome germline SNV calling — can be brought from many hours on multi-core CPU to one minute on a single GPU using hardware that is widely available today. Modern compute hardware offers significant performance that existing bioinformatics tools leave on the table; cuBayes demonstrates how to recover a substantial portion of it for one important analysis. Achieving this without altering the stock version’s algorithmic behavior was a deliberate design objective. We preserved FreeBayes’ mathematical model exactly, so that cuBayes can replace the reference implementation within validated workflows without revalidation. And we structured the implementation around a modular atom/molecule architecture so that the supporting infrastructure — BAM access, decompression, partitioning, pileup construction, batch coordination — is reusable rather than tied to a single algorithm. The persistent gap between hardware capability and bioinformatics software performance is not a technical inevitability, and the present work demonstrates a path to closing it.

### Rapid sample-to-variant turnaround enables new clinical opportunities

Modern sequencing platforms such as the Illumina NovaSeq complete a 60× whole human genome run in approximately 25 hours (Illumina NovaSeq, S1 flow cell). Hardware-accelerated basecalling and demultiplexing — for instance on-instrument DRAGEN BCL Convert — produce FASTQ files in approximately 1.5–2 hours per flow cell under default settings (Illumina, NovaSeq X DRAGEN v4.3.16 BCL Convert release notes, 2025). GPU-accelerated alignment via Parabricks-BWA processes a 60× dataset in under 2 hours^11^. With cuBayes completing germline SNV calling in under one minute on the same class of hardware, a complete sequence-to-variant workflow on a single 60× human genome can now be performed in under 48 hours on a single workstation, end-to-end. This is transformative for clinical settings where turnaround was previously measured in weeks. Glioblastoma carries a median survival of 12–15 months^18^; current precision oncology programs report molecular profiling turnaround on the order of 28 days^19^, leaving little room for tumor board review, therapy selection, and drug acquisition within the actionable window. Compressing analysis to hours rather than weeks makes it feasible to design precision oncology trials and rapid diagnostic programs around comprehensive genomic profiling, even for aggressive cancers and acute presentations of rare disease.

### Faster genomic analysis enables better genomic analysis

The benefit of sub-minute analysis is not limited to clinical settings where speed is intrinsically required. Most genomic analyses in practice are not single-pass automated computations but iterative explorations: an analyst formulates a hypothesis, runs an analysis, inspects the result, identifies an artifact or refinement, adjusts parameters, and re-runs. When each iteration takes hours or days, the cost of re-analysis suppresses the number of hypotheses tested, and human-in-the-loop interpretation becomes serial rather than interactive. When iteration takes minutes, the same analyst can test a meaningfully larger space of hypotheses within a single working session. The same shift benefits method developers. Comprehensive method development requires testing across realistic data scales, but downstream analysis runtimes have historically forced developers to validate on downsampled or single-chromosome subsets during iteration, with full-dataset validation deferred to the end of development^5^. Sub-minute whole-genome runtimes restore full-dataset testing as a routine part of the development loop, reducing the risk that method behavior diverges between development and deployment.

### Drop-in semantic equivalence lowers the cost of adoption

A common pattern in academic GPU acceleration is to introduce alternative algorithms that approximate or replace established methods. Such alternatives require independent validation before they can be substituted into validated workflows — a barrier that is particularly costly in clinical and translational contexts. By preserving FreeBayes’ mathematical model exactly, cuBayes can substitute for the reference implementation within existing pipelines without re-validating downstream analyses. The concordance results we report, with essentially complete agreement at QUAL ≥ 20, support this substitutability in practice. We view this approach — accelerating widely used algorithms while preserving their semantics — as a more direct path to community adoption than developing accelerated alternatives that diverge from established methods.

### The atom/molecule architecture enables sustained acceleration of the broader ecosystem

A single accelerated variant caller does not by itself answer the broader gap in genomic software performance. We designed the atom/molecule architecture with two longer-term goals in mind. First, performance and portability improvements made at the atom level propagate automatically to all consumers: an optimization to BGZF decompression accelerates every molecule and tool that depends on it without per-tool engineering. By the same mechanism, an AMD ROCm backend would extend cross-platform support to all downstream molecules and tools without requiring them to be rewritten. Second, the architecture supports two distinct modes of pipeline integration. In the conventional mode, a complete tool such as cuBayes is placed within a larger workflow alongside other tools, with workflow managers coordinating tool-to-tool data flow through intermediate files. Alternatively, an entire workflow can be expressed at the molecule level, with what would conventionally be tool boundaries instead becoming molecule chains within a single composed application. The latter mode exposes optimization opportunities unavailable to file-based pipelines: shared in-memory buffers across stages, intermediate results passed without materializing to disk, and overlapping computation with I/O across stage boundaries. cuBayes itself uses fine-grain composition internally, sharing partition state between molecules and emitting genotype calls without writing per-region intermediate files.

### Current scope and forward directions

cuBayes in its current form addresses germline SNV calling on aligned short-read data. Several capabilities natively supported by reference FreeBayes are not yet implemented. Indel detection requires variable-length sequence operations that are best handled with different GPU data structures than the fixed-position calls supported here, and is the most prominent capability we plan to add. Haplotype-aware joint genotyping across multiple samples — central to the multi-sample tumor evolution analyses motivating much of the FreeBayes user base — is similarly planned. The current implementation targets NVIDIA CUDA only; an AMD ROCm backend is planned to extend hardware portability. Beyond cuBayes itself, immediate next steps include packaging the workflow within a Nextflow pipeline for production deployment, exposing the GPU-accelerated primitives through Python bindings for use within Jupyter notebooks for interactive analysis, and packaging the atom and molecule libraries — together with the conventions for composing them — as an open software development kit so that other method developers can apply this approach to additional algorithms without reimplementing low-level infrastructure. Source code will be released under a permissive open-source license at the time of journal publication.

### Reproduction and distribution

cuBayes is currently available as a precompiled container image at https://hub.docker.com/r/cubayes/cubayes, with a matching Apptainer/Singularity definition for HPC environments. For reproducible evaluation, we provide a turnkey deployment script https://github.com/shadowfaxbio/cubayes-demo that, on a fresh AWS GPU instance (recommended: g6e.12xlarge), pulls the container image, provisions local NVMe storage, downloads the GIAB benchmark BAMs and the GRCh38 reference FASTA, and executes the cuBayes benchmark end-to-end. Source code will be released under a permissive open-source license at the time of journal publication.

## Methods

### Input data

We used two GIAB benchmark samples for all experiments: HG002 (NA24385, Ashkenazi son) and HG004 (NA24143, Ashkenazi mother), sequenced as 2×250 bp Illumina paired-end reads at approximately 60× coverage. Aligned BAM files were obtained from the GIAB data release, with novoalign V3.02.07 alignments against the GRCh38 reference. Variant calling for both cubayes and reference FreeBayes used the GRCh38 full analysis set with decoy and HLA sequences (GRCh38_full_analysis_set_plus_decoy_hla.fa).

### Hardware and software environment

All benchmarks were performed on a workstation equipped with an AMD Ryzen Threadripper PRO 7975WX processor (32 physical cores), 512 GiB of system memory, and an NVIDIA RTX 6000 Ada Generation GPU (48 GB VRAM, compute capability sm_89). The workstation ran Rocky Linux 9.7 with NVIDIA driver 595.71.05 and CUDA toolkit 12.9. Input BAM files (approximately 130 GB for HG002) resided on a local NVMe-class storage volume.

cuBayes is implemented in CUDA C++20 and was built with nvcc from CUDA toolkit 12.9 (release V12.9.86) at -O3 optimization. The benchmark binary cubayes_actor corresponds to git commit 89f675e (version string v0.2.0-4-g89f675e) on the development branch. Runtime dependencies are quickBAM^14^ v1.0.3, nvcomp 3.0.5, oneAPI TBB 2021.13.0, and libdeflate v1.21.

### Algorithm scope and reference implementation

cuBayes implements germline single-nucleotide variant (SNV) calling against the FreeBayes v1.2.0 algorithm. The current implementation handles diploid single-sample SNV calling on aligned short-read data. Indel detection, multi-nucleotide polymorphism construction, haplotype-aware joint genotyping, and multi-sample analysis are not implemented in this version.

To enable a like-for-like comparison between cubayes and reference FreeBayes, we patched FreeBayes v1.2.0 to disable complex-allele construction (the clumpAlleles() call in AlleleParser.cpp). FreeBayes performs this complex-allele construction unconditionally as part of its internal allele-building procedure and exposes no command-line flag to disable it (--no-mnp filters output but does not skip the construction). Disabling clumpAlleles() brings reference FreeBayes’ work into scope-equivalence with cubayes; without this patch, reference FreeBayes would perform additional work beyond what cubayes implements, which would artificially inflate the measured speedup. The patched FreeBayes is used as the reference for both runtime benchmarking and concordance analysis throughout this work.

The following implementation differences exist between cubayes and reference FreeBayes within their shared SNV-calling scope. (1) cuBayes uses 32-bit single-precision floating point for likelihood and posterior calculations, while reference FreeBayes uses 80-bit extended precision (long double); this affects the magnitude of posterior values but does not affect the genotype ranking that determines calls. (2) cuBayes quantizes combined base- and mapping-quality scores to integer Phred values prior to per-position aggregation, introducing at most ±0.5 Phred per observation; reference FreeBayes retains full per-read precision. (3) cuBayes performs probability arithmetic in log_10_ space; reference FreeBayes uses natural log. These represent the only deliberate algorithmic differences in the SNV path; all default thresholds, priors, and model parameters (theta = 0.01, reference-data factor = 0.9, minimum mapping quality = 1, minimum base quality = 0, mapping-quality adjustment enabled) match FreeBayes defaults and are hard-coded as compile-time constants in this version.

To bound peak memory consumption in extreme-depth regions (centromeres, satellite repeats), cuBayes caps per-region read consumption at 50,000 reads. Regions exceeding this cap are flagged and processed with the available reads. In the data presented here, this cap affects fewer than 100 regions per genome and has no measurable impact on concordance within accessible regions.

### Reference FreeBayes execution

Reference FreeBayes was invoked with the following command line for each genomic region:

~~~
freebayes -f $REF -b $BAM \
   --min-mapping-quality 1 --min-base-quality 0 \
   --theta 0.01 --ploidy 2 \
   --no-indels --haplotype-length 0
~~~

The --no-indels and --haplotype-length 0 flags restrict FreeBayes to single-position SNV calling, matching cubayes’ implemented scope. The default values for --theta, --min-mapping-quality, and -- min-base-quality were specified explicitly to match the constants used by cubayes.

For multi-threaded execution, the genome was partitioned into non-overlapping regions distributed across worker processes using GNU parallel:

~~~
cat $REGIONS | parallel -k -j $NCPUS \
     freebayes <args> --region {} \
   | vcffirstheader \
   | vcfstreamsort -w 1000 \
   | vcfuniq > $OUTPUT
~~~

We benchmarked this orchestration at 12, 32, and 64 worker processes ($NCPUS), corresponding to a configuration matching the 12-thread baseline commonly used in HPC workflows, the workstation’s physical core count, and the workstation’s logical thread count under hyperthreading. The output pipeline (vcffirstheader | vcfstreamsort | vcfuniq) merges per-region VCFs into a single coordinate-sorted output.

### Wall-clock benchmarking

End-to-end wall-clock runtimes were measured with time(1) capturing the complete command invocation, from process start to VCF file closure. For cuBayes runs, the timed command was cubayes_actor <bam> <ref.fa> > <output.vcf>. For reference FreeBayes runs, the timed command was the complete GNU parallel pipeline shown above. All measurements reported in this work are from runs in which the input BAM was already resident on the local storage and the GPU was in its default power state at the start of execution. Each configuration was measured once; we did not perform replicate runs.

### Concordance analysis

Concordance between cubayes and reference FreeBayes was assessed using a custom Python script that compares two VCF files position by position. The metric is sensitive to genotype rather than only to the presence of a call: at each position, the called genotype was extracted as a sorted tuple of allele values (so that 0/1 and 1/0 collapse to the same genotype), and a site was considered concordant when both implementations reported a variant call at the same position with the same genotype. Indel-containing records (REF/ALT length mismatches) were excluded from the comparison, consistent with the SNV-only scope; equal-length multi-nucleotide records were decomposed into per-position SNV records before comparison.

For each tier of analysis, we computed the union of all variant sites called by either implementation, then partitioned the union into four classes: (i) concordant — both implementations called the same variant with matching genotype; (ii) shared sites with discordant genotypes — both implementations called a variant at the position but with different genotypes; (iii) cubayes-only — sites called by cubayes but not reference FreeBayes; (iv) reference-FreeBayes-only — sites called by reference FreeBayes but not cubayes. The reported concordance is the number of concordant sites divided by the size of the union.

Three filter tiers were applied progressively. **Tier 1** is the unfiltered union of all calls from both implementations. **Tier 2** restricts to chr1–22, X, Y and applies QUAL ≥ 20 to both call sets. **Tier 3** further intersects the Tier 2 set with the 1000 Genomes Project pilot accessibility mask^9^. T2 and T3 filters are commonly applied in variant calling pipelines.

## Notes

### Competing Interest Statement

The authors have declared no competing interest.

## References

1. Saunders CJ, Miller NA, Soden SE, Dinwiddie DL, Noll A, Alnadi NA, Andraws N, Patterson ML, Krivohlavek LA, Fellis J, Humphray S, Saffrey P, Kingsbury Z, Weir JC, Betley J, Grocock RJ, Margulies EH, Farrow EG, Artman M, Safina NP, Petrikin JE, Hall KP, Kingsmore SF. Rapid whole-genome sequencing for genetic disease diagnosis in neonatal intensive care units. Sci Transl Med. American Association for the Advancement of Science (AAAS); 2012 Oct 3;4(154):154ra135. PMCID: PMC4283791

2. Clark MM, Hildreth A, Batalov S, Ding Y, Chowdhury S, Watkins K, Ellsworth K, Camp B, Kint CI, Yacoubian C, Farnaes L, Bainbridge MN, Beebe C, Braun JJA, Bray M, Carroll J, Cakici JA, Caylor SA, Clarke C, Creed MP, Friedman J, Frith A, Gain R, Gaughran M, George S, Gilmer S, Gleeson J, Gore J, Grunenwald H, Hovey RL, Janes ML, Lin K, McDonagh PD, McBride K, Mulrooney P, Nahas S, Oh D, Oriol A, Puckett L, Rady Z, Reese MG, Ryu J, Salz L, Sanford E, Stewart L, Sweeney N, Tokita M, Van Der Kraan L, White S, Wigby K, Williams B, Wong T, Wright MS, Yamada C, Schols P, Reynders J, Hall K, Dimmock D, Veeraraghavan N, Defay T, Kingsmore SF. Diagnosis of genetic diseases in seriously ill children by rapid whole-genome sequencing and automated phenotyping and interpretation. Sci Transl Med. American Association for the Advancement of Science (AAAS); 2019 Apr 24;11(489):eaat6177. PMCID: PMC9512059

3. Gorzynski JE, Goenka SD, Shafin K, Jensen TD, Fisk DG, Grove ME, Spiteri E, Pesout T, Monlong J, Baid G, Bernstein JA, Ceresnak S, Chang PC, Christle JW, Chubb H, Dalton KP, Dunn K, Garalde DR, Guillory J, Knowles JW, Kolesnikov A, Ma M, Moscarello T, Nattestad M, Perez M, Ruzhnikov MRZ, Samadi M, Setia A, Wright C, Wusthoff CJ, Xiong K, Zhu T, Jain M, Sedlazeck FJ, Carroll A, Paten B, Ashley EA. Ultrarapid nanopore genome sequencing in a critical care setting. N Engl J Med. Massachusetts Medical Society; 2022 Feb 17;386(7):700–702. PMID: 35020984

4. Wojcik MH, Larkin K, Cipicchio M, Doupnik A, Zhao C, Cech C, Lopez D, Chandrasekar J, Leadbetter J, Mannion J, Berg K, Golkaram M, Osentowski M, Freer M, Lehmann T, Lee WM, Ormbrek E, Prindle MJ, Nabavi M, Chaturvedi A, Seberino C, Baker DN, Williams C, Toledo D, Malolepsza E, Fleharty M, Oza A, Low S, Beggs AH, Genetti CA, Strickland G, Anderson KN, Chung WK, Rehm HL, Hofherr S, Kokoris M, Lennon N, Broad-BCH-Roche Rapid WGS Team. Toward same-day genome sequencing in the critical care setting. N Engl J Med. Massachusetts Medical Society; 2025 Nov 20;393(20):2063–2065. PMCID: PMC12854144

5. Berger B, Yu YW. Navigating bottlenecks and trade-offs in genomic data analysis. Nat Rev Genet. Springer Science and Business Media LLC; 2023 Apr;24(4):235–250. PMCID: PMC10204111

6. Li H. Aligning sequence reads, clone sequences and assembly contigs with BWA-MEM [Internet]. arXiv [q-bio.GN]. 2013. Available from: 10.48550/arXiv.1303.3997

7. McKenna A, Hanna M, Banks E, Sivachenko A, Cibulskis K, Kernytsky A, Garimella K, Altshuler D, Gabriel S, Daly M, DePristo MA. The Genome Analysis Toolkit: a MapReduce framework for analyzing next-generation DNA sequencing data. Genome Res. Cold Spring Harbor Laboratory; 2010 Sep;20(9):1297–1303. PMCID: PMC2928508

8. Garrison E, Marth G. Haplotype-based variant detection from short-read sequencing [Internet]. arXiv [q-bio.GN]. 2012. Available from: 10.48550/arXiv.1207.3907

9. 1000 Genomes Project Consortium, Auton A, Brooks LD, Durbin RM, Garrison EP, Kang HM, Korbel JO, Marchini JL, McCarthy S, McVean GA, Abecasis GR. A global reference for human genetic variation. Nature. Springer Science and Business Media LLC; 2015 Oct 1;526(7571):68–74. PMCID: PMC4750478

10. Heldenbrand JR, Baheti S, Bockol MA, Drucker TM, Hart SN, Hudson ME, Iyer RK, Kalmbach MT, Kendig KI, Klee EW, Mattson NR, Wieben ED, Wiepert M, Wildman DE, Mainzer LS. Recommendations for performance optimizations when using GATK3.8 and GATK4. BMC Bioinformatics. Springer Science and Business Media LLC; 2019 Nov 8;20(1):557. PMCID: PMC6842142

11. Marangoni S, Furia F, Charrance D, Fant A, Di Dio S, Trova S, Spirito G, Musacchia F, Coppe A, Gustincich S, Vecchi M, Landuzzi F, Cavalli A. GeNePi: a graphics processing unit enhanced next-generation bioinformatics pipeline for whole-genome sequencing analysis. Brief Bioinform. Oxford University Press (OUP); 2026 Jan 7;27(1):bbag001. PMCID: PMC12832024

12. Ren S, Bertels K, Al-Ars Z. Efficient acceleration of the pair-HMMs forward algorithm for GATK HaplotypeCaller on graphics processing units. Evol Bioinform Online. SAGE Publications; 2018 Mar 12;14:1176934318760543. PMCID: PMC5858735

13. Poplin R, Chang PC, Alexander D, Schwartz S, Colthurst T, Ku A, Newburger D, Dijamco J, Nguyen N, Afshar PT, Gross SS, Dorfman L, McLean CY, DePristo MA. A universal SNP and small-indel variant caller using deep neural networks. Nat Biotechnol. Springer Science and Business Media LLC; 2018 Nov;36(10):983–987. PMID: 30247488

14. Pitman A, Huang X, Marth GT, Qiao Y. quickBAM: a parallelized BAM file access API for high-throughput sequence analysis informatics. Bioinformatics [Internet]. Oxford University Press (OUP); 2023 Aug 1;39(8). Available from: 10.1093/bioinformatics/btad463 PMCID: PMC10412403

15. Qiao Y, Quinlan AR, Jazaeri AA, Verhaak RG, Wheeler DA, Marth GT. SubcloneSeeker: a computational framework for reconstructing tumor clone structure for cancer variant interpretation and prioritization. Genome Biol. Springer Science and Business Media LLC; 2014 Aug 26;15(8):443. PMCID: PMC4180956

16. giab_data_indexes: This repository contains data indexes from NIST’s Genome in a Bottle project [Internet]. Github; [cited 2026 Jun 12]. Available from: https://github.com/genome-in-a-bottle/giab_data_indexes

17. Krusche P, Trigg L, Boutros PC, Mason CE, De La Vega FM, Moore BL, Gonzalez-Porta M, Eberle MA, Tezak Z, Lababidi S, Truty R, Asimenos G, Funke B, Fleharty M, Chapman BA, Salit M, Zook JM, Global Alliance for Genomics and Health Benchmarking Team. Best practices for benchmarking germline small-variant calls in human genomes. Nat Biotechnol. Springer Science and Business Media LLC; 2019 May;37(5):555–560. PMCID: PMC6699627

18. Stupp R, Mason WP, van den Bent MJ, Weller M, Fisher B, Taphoorn MJB, Belanger K, Brandes AA, Marosi C, Bogdahn U, Curschmann J, Janzer RC, Ludwin SK, Gorlia T, Allgeier A, Lacombe D, Cairncross JG, Eisenhauer E, Mirimanoff RO, European Organisation for Research and Treatment of Cancer Brain Tumor and Radiotherapy Groups, National Cancer Institute of Canada Clinical Trials Group. Radiotherapy plus concomitant and adjuvant temozolomide for glioblastoma. N Engl J Med. New England Journal of Medicine (NEJM/MMS); 2005 Mar 10;352(10):987–996. PMID: 15758009

19. Worst BC, van Tilburg CM, Balasubramanian GP, Fiesel P, Witt R, Freitag A, Boudalil M, Previti C, Wolf S, Schmidt S, Chotewutmontri S, Bewerunge-Hudler M, Schick M, Schlesner M, Hutter B, Taylor L, Borst T, Sutter C, Bartram CR, Milde T, Pfaff E, Kulozik AE, von Stackelberg A, Meisel R, Borkhardt A, Reinhardt D, Klusmann JH, Fleischhack G, Tippelt S, Dirksen U, Jürgens H, Kramm CM, von Bueren AO,Westermann F, Fischer M, Burkhardt B, Wößmann W, Nathrath M, Bielack SS, Frühwald MC, Fulda S, Klingebiel T, Koscielniak E, Schwab M, Tremmel R, Driever PH, Schulte JH, Brors B, von Deimling A, Lichter P, Eggert A, Capper D, Pfister SM, Jones DTW, Witt O. Next-generation personalised medicine for high-risk paediatric cancer patients -The INFORM pilot study. Eur J Cancer. Elsevier BV; 2016 Sep;65:91–101. PMID: 27479119

